# *Lactococcus lactis* subsp. *cremoris* C60 creates a type I regulatory T cell-dominant anti-inflammatory intestinal environment through functional modification of dendritic cells

**DOI:** 10.1101/2024.12.03.626564

**Authors:** Suguru Saito, Nanae Kakizaki, Alato Okuno, Toshio Maekawa, Noriko M. Tsuji

**Author notes:** Correspondence Suguru Saito, Ph.D; Noriko M. Tsuji, Ph.D., (ext 2261).

## Abstract

Lactic acid bacteria (LAB)-mediated probiotics have the capability to modulate intestinal immunity, creating an anti-inflammatory environment. Among the immune-modulatory effects, interleukin (IL)-10-producing CD4^+^ T cells are often discussed for their anti-inflammatory roles. While regulatory T cells (Tregs) are a well-known subset, type I regulatory T (Tr1) cells, which also produce abundant IL-10, receive less attention in LAB-induced anti-inflammatory effects. Here, we report that *Lactococcus lactis* subsp. *cremoris* C60, a probiotic LAB strain, increases IL-10-producing Tr1 cells in the intestinal environment, contributing to resistance against inflammatory conditions such as dextran sodium sulfate (DSS)-induced colitis. Intragastric administration of heat-killed (HK)-C60 significantly increased IL-10^+^CD4^+^ T cells in intestinal tissues compared to control treatment. These populations predominantly consisted of Forkhead box protein P3 (Foxp3)-negative cells, indicating that Tr1 cells were primarily expanded by HK-C60 stimulation. Further analysis revealed that IL-10^+^ Tr1 cells could be classified into two major subpopulations: IL-10 single-positive (SP) and IFN-γ/IL-10 double-positive (DP) cells. An *in vitro* co-culture system using CD4^+^ T cells and bone marrow-derived dendritic cells (BMDCs) demonstrated that HK-C60 stimulation modified dendritic cell function, promoting Tr1 cell differentiation. Moreover, pro-inflammatory IFN-γ^+^CD4^+^ T (Th1) cells differentiated into IL-10^+^ Tr1 cells in the presence of HK-C60-stimulated BMDCs, with the effect being particularly pronounced in DP Tr1 cells. HK-C60-administration attenuated inflammation and reduced pro-inflammatory leukocyte infiltration in the colon of DSS-induced colitis mice, accompanied by the expansion of DP Tr1 cells. The suppressive effect of intestinal CD4^+^ T cells in HK-C60-treated mice was IL-10-dependent, as demonstrated by an *in vitro* IL-10 neutralizing assay. Thus, C60-based probiotics promote the creation of an anti-inflammatory intestinal environment by increasing IL-10^+^ Tr1 cells rather than Tregs.

## Introduction

Probiotics utilizing lactic acid bacteria (LAB) are a well-known approach for improving the intestinal environment [1]. A key effect of probiotic LAB is frequently attributed to the functional modulation of immune cells, particularly myeloid cells [2, 3]. Similar to the recognition of pathogenic bacteria, LAB are primarily recognized by extracellular pattern recognition receptors (PRRs) and subsequently internalized into the cytosol [4, 5]. This series of responses stimulates myeloid cells, leading to enhanced functional activity and alterations in various immune responses [6, 7]. Such functional changes are often transferred to adaptive immunity, exemplified by T cell modulation, ultimately resulting in the comprehensive conditioning of immune functions [8,9].

*Lactococcus lactis* subsp. *cremoris* C60 is a probiotic strain with demonstrated immunomodulatory effects on myeloid cells [7, 10]. It stimulates antigen-presenting cells (APCs), such as dendritic cells (DCs) and macrophages, resulting in increased cytokine production and enhanced antigen presentation [10, 11]. The immunomodulatory effects of C60 extend to both CD4^+^ and CD8^+^ T cells, suggesting that this strain universally enhances the antigen presentation machinery in APCs, regardless of MHC class restriction [10, 11, 12]. However, in the intestinal environment, C60’s effects are more pronounced on CD4^+^ T cell functions. Previous studies have shown that intragastric (i.g.) administration of C60 increases IFN-γ^+^CD4^+^ T (Th1) cells in the intestinal tissue of mice through enhanced APC activity [10, 11]. Furthermore, C60 administration restored the Th1 cell population via functional upregulation of DCs in interleukin-18 knockout (IL-18 KO) mice, which exhibit severe deficiencies in effector CD4^+^ T cells in the intestine [10]. This finding suggests that C60-induced functional modifications in APCs operate through diverse mechanisms, independent of canonical pro-inflammatory cytokines like IL-18, which is typically indispensable for Th1 cell differentiation [13]. Maintaining intestinal homeostasis requires a delicate balance between pro-inflammatory and anti-inflammatory immune responses [14]. In probiotic studies, including those involving C60, the upregulation of Th1 cells has been extensively examined [15]. However, the concurrent effects of probiotics on the generation of IL-10-producing CD4^+^ T cells have received comparatively little attention. Understanding how a probiotic strain influences both pro-inflammatory and anti-inflammatory immune responses is crucial for evaluating its role in maintaining immunological balance.

Given this, the present study investigates the effects of C60 on anti-inflammatory T cell generation in the intestinal environment. Administration of heat-killed (HK) C60 significantly increased IL-10-producing CD4^+^ T cells alongside Th1 cells in intestinal tissues, such as Peyer’s patches, and the lamina propria of the small (sLP) and large intestines (cLP). Interestingly, C60 predominantly increased forkhead box P3 (Foxp3)-negative CD4^+^ T cells within the IL-10-producing population rather than Foxp3^+^CD4^+^ T cells, which are commonly associated with regulatory T cells (Tregs) [16]. Foxp3^-^negative CD4^+^ T cells, classified as type 1 regulatory T (Tr1) cells, are known to produce abundant IL-10 and play a critical role in suppressing inflammation [17]. Further, we found that C60-stimulated DCs enhanced IL-10 production and upregulated antigen-presenting molecules, resulting in increased IL-10^+^ Tr1 cell generation. Interestingly, these DCs induced the differentiation of IL-10^+^ Tr1 cells from pro-inflammatory Th1 cells, with a particularly pronounced effect on the differentiation of IFN-γ^+^IL-10^+^ Tr1 cells. By promoting these immunological changes, HK-C60 administration alleviated symptoms of dextran sodium sulfate (DSS)-induced colitis in mice. This effect was accompanied by an increase in intestinal IL-10^+^CD4^+^ T cells, predominantly composed of Tr1 cells, alongside a reduction in inflammatory myeloid cells and T cells.

Thus, C60 is a unique probiotic strain that influences both pro-inflammatory and anti-inflammatory T cell generation through functional modifications of DCs, offering potential therapeutic benefits for intestinal inflammation.

## Materials and methods

### Mice

BALB/c wild-type (WT) mice were purchased from Japan Clea (Tokyo, Japan). IL-10-green fluorescent protein (GFP) reporter mice (B6.129S6-*Il10^tm1Flv^*/J) and DO11.10 mice (C.Cg-Tg(DO11.10)10Dlo/J) were purchased from the Jackson laboratory (Bar harbor, MA, USA). All mice were bred in-house and maintained in a specific pathogen-free (SPF) condition with 12 h day/night cycles and were allowed free access to food and water. To maintain similar microbiota and intestinal environment, all the mice used in this study had been bred in the same facility at least for 3 months. Gender-matched adult (8-16 weeks) mice were used for each experiment. To investigate the immunomodulatory effect of LAB under physiological condition, the mice received intragastric (i.g.) administration of saline (200 μl) or heat-killed (HK)-C60 (200 μl of 5.0×10^9^ CFU/mL in saline) every day for 14 days. After completion of the administration schedule, the mice were sacrificed and used for analysis. All the animal experiment protocols were approved by the animal care and use committee of AIST (No. 109).

### Lactic acid bacteria culture

*Lactococcus lactis* subsp. *Cremoris* C60 was cultured by following the method described in a previous report [10]. Briefly, the bacteria were cultured in MRS broth (BD bioscience, Franklin Lakes, NJ, USA) at 37 ℃ for 24h. The bacterial colony forming unit (CFU/mL) was calculated in all cultures. For HK-C60 preparation, the bacteria were heated at 95°C for 10 min, then the bacterial cells were precipitated by centrifugation at 10,000 *g* for 5 min. After washing with PBS, the cell pellet was resuspended in 0.9% NaCl (saline) or phosphate-buffered saline (PBS). The sample was used as Heat-killed (HK)-C60.

### Reagents, antibodies and cell isolation kits

Phorbol 12-myristate 13-acetate (PMA) were ionomycin were purchased from Sigma Aldrich (St. Louis, MO, USA). Collagenase-D, collagenase type-I, DNase and Dynabeads™ Mouse T-Activator CD3/CD28 for T-Cell expansion and activation were all purchased from Thermo Fisher Scientific (Waltham, MA, USA). A Cytofix/Cytoperm kit with GolgiStop^TM^ and thioglycolate broth was purchased from BD BioscienceFOXP3 Fix/Perm Buffer Set, anti-CD3 (17A2), anti-CD4 (GK1.5), anti-CD8 (53-6.7), anti-CD45 (30-F11), anti-CD11b (M1/70), anti-F4/80 (BM8), anti-CD11c (N418), anti-CD80 (16-10A1), anti-CD86 (GL-1), anti-I-A/I-E (M5/114.15.2), anti-IFN-γ (XMG1.2), anti-IL-4 (11B11), anti-IL-10 (JES5-16E3), anti-IL-17A (TC11-18H10.1), anti-TNF (MP6-XT22), anti-CD16/CD32 (93) and Zombie Aqua™ Fixable Viability Kit were purchased from BioLegend (San Diego, CA, USA). Recombinant murine granulocyte macrophage-colony stimulating factor (rmGM-CSF) was purchased from peprotech (Cranbury, NJ, USA). Lipopolysaccharide (LPS) was purchased from Invivogen (San Diego, CA, USA). Dextran sodium sulfate was purchased from Iwai Chemicals Company (Tokyo, Japan). CD11c Micro Beads UltraPure and CD4 ^+^ T Cell Isolation Kit, mouse were purchased from Miltenyi Biotec (Bergisch Gladbach, Germany).

### Flow cytometry

Cell surface markers and intracellular cytokines were analyzed by a flow cytometer (FACS Aria I; BD Biosciences, Franklin Lakes, NJ, USA). The cells were first incubated with Fc receptor blocker (anti-CD16/CD32) at 4°C for 10 min. For surface marker staining, the cells were incubated with the antibody in PBS/2% FBS at 4°C for 30 min. For effector T cells characterization by following the cytokine production, the samples were stimulated prior to intracellular staining. For IFN-γ, IL-4 or IL-17A detection, the samples were treated with PMA (100 ng/mL) and ionomycin (250 ng/mL) in the presence of GolgiStop^TM^ (1l/mL) at 37°C for 6 h. For IL-10 detection, the samples were re-stimulated with anti-CD3 mAb (10 µg/mL) and anti-CD28 mAb (2 μg/mL) at 37°C for 48 h. At the last 6 h, the samples were re-simulated with PMA (100 ng/mL) and ionomycin (250 ng/mL) in the presence of GolgiStop^TM^ (1 μl/mL). The sample was firstly stained with antibodies for extracellular markers, then were treated with BD Cytofix/CytoPerm buffer at 4°C for 20 min. After being washed with BD Perm/Wash buffer, the cells were incubated with the antibody for intracellular cytokine staining at 4°C for 30-60 min. The dead cells were excluded by Live/Dead staining in each analysis. All data were analyzed by BD FACS Diva (BD Bioscience) or FlowJo (BD Bioscience).

### Preparation of Peyer’s patch cells

Peyer’s patch (PP)-derived cells were isolated by following the protocol described in a previous report [10]. Briefly, the excised small intestinal PPs were incubated in washing buffer (PBS containing 10% FBS, 100 U/mL penicillin, 100 U/mL streptomycin, 20 mM EDTA and 100 mM DTT) at 37°C for 20 min with stirring. The PPs were collected by decantation onto a 70 µM cell strainer and then briefly washed with cell culture medium (RPMI 1640 supplemented with 10% fetal bovine serum (FBS), 50 μM 2-mercaptoethanol (2-ME), 10 mM HEPES, 100 U/mL penicillin and 100 mg/mL streptomycin). The PPs were transferred to digestion buffer (RPMI1640 containing 10% FBS, 1 mg/mL collagenase type-I, 50 ug/mL DNase) and incubated at 37°C for 30 min with stirring. After digestion, the PPs were mechanically crushed on a 70 µM cell strainer, then the cells were washed with cell culture medium. The sample was filtered through a 40 µM cell strainer again, then then were collected by centrifugation at 300 *g* for 5 min. the precipitated cells were used as PP isolate cells.

### Preparation of lamina propria leukocytes

Lamina propria (LP) cells were isolated by following the protocol described in a previous report [10]. Briefly, small intestine or colon was isolated from mouse and was washed thoroughly in ice-cold PBS. The intestine was cut to small pieces (5-10 mm) and incubated in a washing buffer (PBS containing 10% FBS, 10 mM EDTA, 20 mM HEPES, 100 U/mL penicillin, 100 U/mL streptomycin, 2 mM sodium pyruvate) at 37°C for 30 min with stirring. After incubation, the tissue pieces were washed in PBS several times and then further cut to small pieces (1-2 mm) in digestion buffer (RPMI 1640 medium containing 10% FBS, 20 mM HEPES, 100 U/mL penicillin, 100 U/mL streptomycin, 1 mg/mL collagenase D, 50 μg/mL DNase I). The sample was incubated at 37°C for 30 min with stirring. After digestion, the sample was filtered through on a 70 µM cell strainer. Undigested tissue pieces were mechanically crushed on the cell strainer. The cells were washed with cell culture medium, and then filtered through a 40 µM cell strainer. The cells were collected by centrifugation at 300 *g* for 5 min. Finally, the leukocyte population was collected by density gradient using percoll. The leukocytes were collected and washed once with cell culture medium. The precipitated cells were used as isolated leukocytes from small intestinal LP (sLP) or colon LP (cLP), respectively.

### Preparation of bone marrow derived dendritic cells (BMDCs)

BMDCs were prepared by following the protocol described in a previous report [10]. The BM cells were flushed out from the tibia and femur with a 10 mL syringe with a 27G needle containing cell culture medium. The cell suspension was filtered through a 70 µM cell strainer and washed once with cell culture medium. After Red blood cells (RBCs) lysis at RT for 10 min, the cells were collected by centrifugation at 300 *g* for 5 min. BM-derived cells (3.0×10^5^/mL) were seeded in 6 well plate with DC culture medium (cell culture medium supplemented with 20 ng/mL of rmGM-CSF), and the plate was incubated at 37°C for 8 days. The half volume of medium was replaced to the fresh DC culture medium in day 6 and 6. At day 8 of culture, the cells were harvested and CD11c^+^ population was enriched using CD11c Micro Beads UltraPure (Miltenyi Biotec, Bergisch Gladbach, Germany). The quality of BMDCs was analyzed by flow cytometry in every culture. Samples with more than 90% of CD11c^+^ were used for experiment.

### In vitro antigen presentation assay

DO11.10 mice received i.g. administration of HK-C60 (200 μl of 5.0×10^9^ CFU/mL in saline) for 14 days, then splenic CD4^+^ T cells were isolated from small intestinal PP-derived cells using CD4 ^+^ T Cell Isolation Kit, mouse (Miltenyi). The CD4^+^ T cells (5.0×10^6^/mL) and BMDCs (1.0×10^6^/mL) were co-cultured in the presence of OVA_323-339_ peptide (100 ng/mL) and vehicle or HK-C60 (5.0×10^7^ CFU/mL) at 37°C for 72 h. The samples were re-stimulated with PMA (100 ng/mL) and ionomycin (250 ng/mL) in the presence of GolgiStop^TM^ (1 μl/mL) in the last 6 h followed by staining for flow cytometry analysis. The purity of isolated CD4^+^ T cells was assessed by flow cytometry and the samples more than 90% of CD4^+^ were used for experiments.

### In vitro Tr1 cell differentiation assay

Splenic CD4+ T cells were isolated from DO11.10 mice following then method described above. The CD4+ T cells (3.0×10^7^/mL) were first culture with anti-CD3/CD28 microbeads (at the ratio=1:1) in the presence of IFN-γ (10 ng/mL), IL-12 (10 ng/mL) and IL-2 (10 ng/mL) at 37°C for 72 h. The expanded and differentiated CD4^+^ T cell in Th1 cells (IFN-γ+CD4+ T cells) were enriched by flow cytometry sorting. The Th1 cells (5.0×10^6^/mL) were further cultured with or without BMDCs (1.0×10^6^/mL) in the presence of OVA_323-339_ peptide (100 ng/mL) and vehicle or HK-C60 (5.0×10^7^ CFU/mL) at 37°C for 72 h. The samples were re-stimulated with PMA (100 ng/mL) and ionomycin (250 ng/mL) in the presence of GolgiStop^TM^ (1 μl/mL) in the last 6 h followed by staining for flow cytometry analysis.

### Dextran sodium sulfate (DSS)-induced colitis

The mice received saline or HK-C60 for 14 days following protocol described above. At the last 5 days, the mice were supplied with regular water or 5% DSS containing water. Throughout this treatment, the body weight (BW) of each mouse was measured daily. At day 14, the mice were sacrificed and used for analysis. The intestinal tissues were subjected to histological and immunological analyses. The colon section was fixed with 10% buffer formalin and embedded to paraffin for Hematoxylin and Eosin (H&E) staining.

### Preparation of lymph node cells

Lymph node (LN)-derived cells were prepared from mesenteric lymph nodes (MLNs). Briefly, the extracted MLNs were mechanically crushed on a 70 µM cell strainer and washed with cell culture medium. The cells were then centrifuged at 300 *g* for 5 min. After being washed once with cell culture medium, the cells were finally collected by centrifugation at 300 *g* for 5min. The precipitated cells were resuspended and used as LN-derived cells. CD4^+^ T cells were isolated from the LN-derived cells following protocol described above.

### Preparation of thioglycolate elucidated peritoneal macrophages (TPMs)

Mice received intraperitoneal (i.p.) injection of 2.5 ml of 3 % thioglycolate. After 84-96 h of the injection, the peritoneal leakage containing leukocytes was collected from peritoneal cavity. To increase the purity of TPMs, the collected cells were seeded in 100 mm cell culture dish with cell culture medium and incubated at 37°C for 2 h. After washing with PBS, the adherent cells were harvested as TPMs. The quality of TPMs was analyzed by flow cytometry in every culture. The sample with more than 90% of CD11b^+^F4/80^+^ was used for experiment.

### In vitro inflammatory suppression assay

CD4^+^ T cells (target cells) were isolated from MLNs of DSS-induced colitis mice. TPMs (1.0×10^7^/mL, target cells) were primed with LPS (100 ng/mL) on 24 well plate at 37°C for 6 h, then the cells were washed with PBS twice. The CD4^+^ T cells (effector cells) were isolated from small intestinal PPs of saline or HK-C60-administrated mice. All CD4^+^ T cells were isolated the samples by using CD4^+^ T cell isolation kit (Miltenyi). The effector CD4^+^ T cells (1.0×10^7^/mL) were stimulated with Dynabeads™ Mouse T-Activator CD3/CD28 (Thermo fisher scientific, at the ratio=1:1) in 24 well plate at 37°C for 72 h. The activated CD4^+^ T cells (1.0×10^7^/mL) were re-seeded in chamber (0.45 μm) which was finally inserted into the well containing target CD4^+^ T cells (1.0×10^7^/mL) with anti-CD3/CD28 microbeads stimulation (at the ratio=1:1) or LPS-primed TPMs (1.0×10^7^/mL). The culture was incubated at 37°C for 16 h. The cytokine production in target CD4^+^ T cells and TPMs was analyzed by flow cytometry, respectively.

### Micro array

The small intestinal Peyer’s patches (PPs) were collected from the mice with oral administration of saline or HK-C60 for 14 days. The CD11c^+^ DCs were isolated from the PP-derived cells using CD11c Micro Beads Ultra Pure (Miltenyi Biotec). The total RNA was isolated from the DCs using TRIzol^TM^ reagent (Thermo Fisher Scientific). RNA purity was evaluated in each sample and the samples with A_260_/A_280_=1.8-2.0 were used for subsequent assay procedure. DNA probes were amplified using Ambion® WT Expression Kit (Thermo Fisher Scientific) from total RNA and hybridized to the Mouse Gene 2.0 ST Array (Affymetrix, Santa Clara, CA, USA).

### Real-time quantitative polymerase chain reaction (real-time qPCR)

Total RNA was isolated from sorted cells by using RNeasy Mini Kit (QIAGEN, Germany). For cDNA synthesis, 500 ng of total RNA was used in RT reaction with PrimeScript RT reagent kit (TaKaRa, Tokyo, Japan). qPCR was performed by using TB Green Premix Ex Taq II reagent (TaKaRa) in Thermal Cycler Dice (TaKaRa). The expression of *Gapdh* mRNA was used as an internal control. The difference of gene expression was quantified by ΔCt method. All procedures were followed by manufacturer’s protocols. The primer sequences used for specific amplification of the respective genes are shown in supplemental table 1.

### Enzyme-linked immunosorbent assay (ELISA)

For cytokine measurement by ELISA, the cultured medium was harvested and stored at −80°C until use. Colon section was chopped by scissors, then digested with collagenase type I (1 mg/mL) at 37°C for 30 min. The samples were filtered through on a 70μm of cell strainer and the flow through was mixed with equal volume of 2x RIPA buffer. The sample was stored at −80℃ until use. The concentration of each cytokine was measured by Duo set ELISA kit (R&D systems, Minneapolis, MN, USA) for each target. All procedure was performed by following product manual.

### Statistics

For comparison between two groups, a student t-test was used to analyze the data for significant differences. For comparison between more than three groups, one-way analysis of variance (ANOVA) was used for significant differences. Values of *p* < 0.05, *p* < 0.01, and *p* < 0.001 were regarded as significant.

## Results

### HK-C60 administration increases IL-10-Producing CD4^+^ T Cells in the Intestinal Environment

Our previous studies demonstrated that C60 is a probiotic strain capable of enhancing the effector functions of CD4^+^ T cells in the intestinal environment [10,11]. However, the scope of our previous screening was limited. To further investigate the effects of C60 on T cell functions, we performed a comprehensive characterization of T-lineage cells based on their cytokine production profiles in intestinal tissues.

Mice were administered saline or heat-killed (HK)-C60 intragastrically (i.g.) daily for 14 days. Intestinal tissues, including Peyer’s patches (PPs), small intestinal lamina propria (sLP), and colonic lamina propria (cLP), were collected and subjected to flow cytometry analysis for T cell characterization (Figure 1A). The frequencies of CD3^+^ pan-T cells were comparable between the saline- and HK-C60-treated groups across all tissues analyzed (Supplemental Figure 1A). Similarly, the frequencies of both CD4^+^ and CD8^+^ T cells were consistent between the two groups in the intestinal tissues (Supplemental Figures 1B, C). Regarding CD4^+^ T cell effector functions, we identified two major cytokine-producing populations: IFN-γ^+^CD4^+^ T (Th1) cells and IL-10^+^CD4^+^ T cells. Both populations showed significantly increased frequencies following HK-C60 administration compared to control treatment in the intestinal tissues. The increased frequency of Th1 cells aligns with our previous findings, reaffirming the reproducible effect of HK-C60 in expanding this effector CD4+ T cell population (Figures 1B, D, E, and F) [10, 11]. Notably, it was a novel observation that HK-C60 administration also increased the frequency of IL-10^+^CD4^+^ T cells in all analyzed intestinal tissues compared to the control group (Figures 1C, D, E, and F). In contrast to the CD4^+^ T cell subsets, HK-C60 administration did not influence the effector functions of CD8^+^ T cells (Supplemental Figures 1D-F).

**Figure 1.**
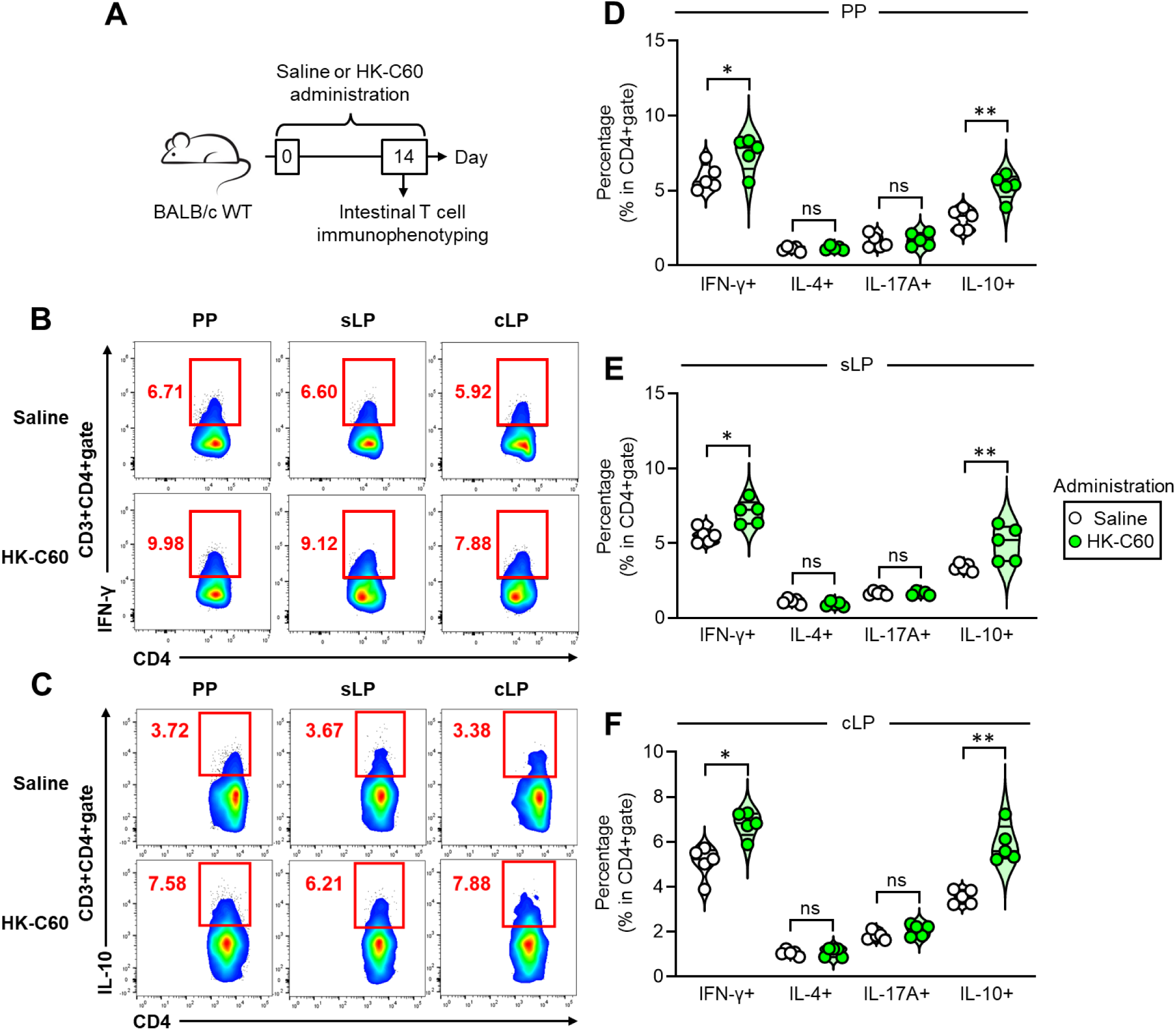
Characterization of intestinal T cells in C60-administered mice. A) Experimental design of intestinal T cell analysis. WT mice received i.g. administration of saline or HK-C60 for 14 days. The leukocytes were isolated from small intestinal PPs, sLP or cLP, and subjected to flow cytometry analysis. B-C) Representative plots of IFN-γ^+^CD4^+^ T cells (B) or IL-10^+^CD4^+^ T cells (C) in each tissue. D-F) Cumulative percentage values of IFN-γ^+^CD4^+^ T (Th1) cells, IL-4^+^CD4^+^ T (Th2) cells, IL-17A^+^CD4^+^ T (Th17) cells and IL-10^+^CD4^+^ T cells in small intestinal PP (D), sLP (E) and cLP (F), respectively. The cumulative data are shown as mean ± SEM values of five samples from two independent experiments. Student t-test was used to analyze data for significance. \**p* < 0.05, \*\**p* < 0.01 and \*\*\**p* < 0.001, ns is not significant.

The IL-10 expression was also validated using the IL-10-GFP reporter system in mice. GFP expression was significantly increased in CD4^+^ T cells derived from the small intestinal PPs of HK-C60-administered mice compared to controls (Supplemental Figure 2A, B). The frequency of IL-10-GFP^+^CD4^+^ T cells positively correlated with the IL-10^+^CD4^+^ T cells, confirming that the direct detection of IL-10 in flow cytometry analysis (Supplemental Figure 2C).

Thus, HK-C60 administration predominantly promotes the expansion of IFN-γ- and IL-10-producing CD4^+^ T cells in the intestinal environment.

### C60 administration increases Tr1 cells in intestinal IL-10-producing CD4^+^ T cells

We next examined the detailed characteristics of IL-10-producing CD4^+^ T cells in the intestinal environment. Peyer’s patch (PP)-derived cells were isolated from mice administered with saline or heat-killed (HK)-C60 for 14 days and analyzed. Given that regulatory T cells (Tregs) are generally considered the major IL-10-producing CD4^+^ T cell subset [16], we first assessed the frequencies of Foxp3^+^ and Foxp3^-^ populations within the IL-10^+^CD4^+^ T cell compartment (Figure 2A).

**Figure 2.**
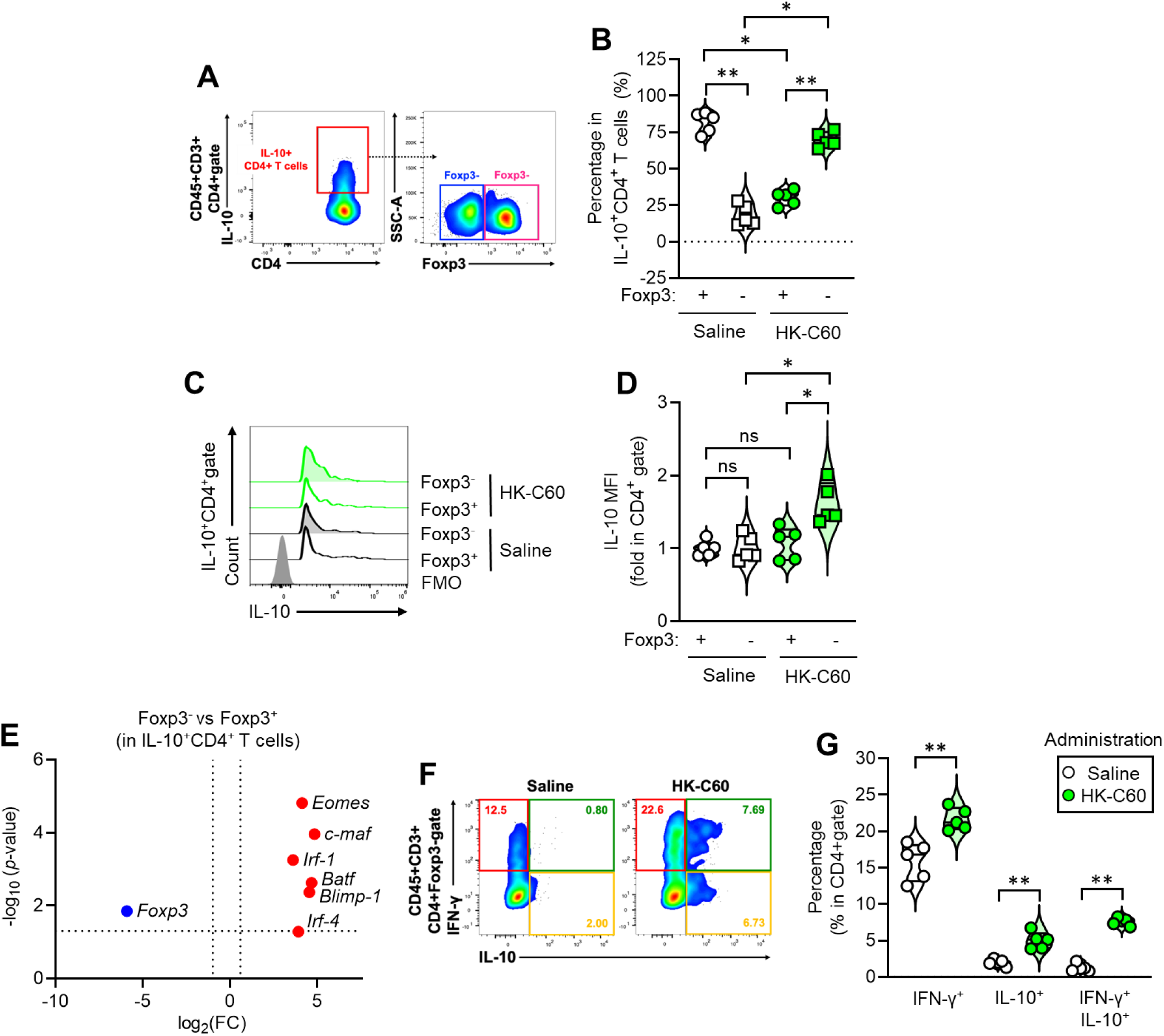
Dominant increase of Foxp3^-^Tr1 cells in intestinal environment of C60-administered mice. The mice received i.g. administration of saline or HK-C60 for 14 days following the schedule described in Figure 1A. The small intestinal PP-derived cells were subjected to analyses. A) Gating strategy for identification of Foxp3^+^ (Treg) and Foxp3^-^ (Tr1 cell) populations in IL-10^+^CD4^+^ T cells. B) Cumulative percentage values of Foxp3^+^ or Foxp3^-^ cells in IL-10^+^CD4^+^ T cell population. C-D) Representative histograms (C) and cumulative MFI values (fold change) (D) of IL-10 productions in Foxp3^+^ or Foxp3^-^IL-10^+^CD4^+^ T cells. E) Gene expression profile of transcription factors. IL-10^+^CD4^+^ T cells were isolated from PP-derived cells of HK-C60-administered mice. The total RNA was subjected to gene expression analysis by real-time PCR. The isolated cells from three mice were pooled and five different batched samples were prepared in Foxp3^+^ and Foxp3^-^ groups, respectively. The difference of gene expression was quantified by ΔCt method. F) Representative plots of subpopulations in Foxp3^-^ CD4^+^ T (Tr1) cells. Each subpopulation was determined by cytokine production patters, such as IFN-γ^+^ (Single Positive; SP), IL-10^+^ (SP), or IFN-γ^+^IL-10^+^ (DP). G) Cumulative percentage values of each subpopulation in Tr1 cells. The cumulative data are shown as mean ± SEM values of five samples from two independent experiments. All MFI values are represented as fold changes (the average IL-10 MFI value of Foxp3^+^ in saline administered group was used for base=1). One-way ANOVA was used to analyze data for significance. \**p* < 0.01 and \*\**p* < 0.001, ns is not significant.

Under basal conditions (saline administration), the Foxp3^+^ population accounted for nearly 80% of the IL-10^+^CD4^+^ T cells, classifying them as Tregs. Conversely, the Foxp3^-^ population was relatively minor, comprising less than 10%. Interestingly, HK-C60 administration induced a notable shift in the ratio of Foxp3^+^ to Foxp3^-^ populations. In HK-C60-treated mice, Foxp3^-^ cells represented over 60% of the IL-10^+^CD4^+^ T cell population, significantly surpassing the Foxp3^+^ population, which constituted approximately 35% (Figure 2B). Moreover, we observed an unexpected change in the intensity of IL-10 expression between Foxp3^+^ and Foxp3^-^ cells. In the basal condition, the mean fluorescence intensities (MFIs) of IL-10 were comparable between Foxp3^+^ and Foxp3^-^ CD4^+^ T cells. However, following HK-C60 administration, the IL-10 MFI was significantly higher in Foxp3^-^ CD4^+^ T cells compared to Foxp3^+^ cells. This finding indicates that Foxp3^-^ cells are not only more abundant but also produce higher levels of IL-10 in the intestinal environment of HK-C60-treated mice (Figures 2C, D).

To further characterize the intestinal IL-10^+^Foxp3^-^ CD4^+^ T cells expanded by HK-C60 administration, we analyzed their gene expression profiles. Foxp3^-^CD4^+^ T cells are typically classified as a different regulatory T cell subset, such as Tr1 cells [17]. However, recent studies suggest that additional transcription factors contribute to Tr1 cell differentiation [17, 18, 19]. To investigate this, we assessed the expression of transcription factors including eomesodermin (Eomes), MAF bZIP transcription factor (c-Maf), basic leucine zipper transcription factor, ATF-like (Batf), B lymphocyte-induced maturation protein-1 (Blimp-1), interferon regulatory factor 1 (Irf-1), and Irf-4, alongside Foxp3. Foxp3^+^ and Foxp3^-^ cells were sorted from small intestinal PP-derived cells of HK-C60-treated mice. Total RNA from these cells was subjected to real-time qPCR for gene expression analysis. Interestingly, Foxp3^-^IL-10^+^CD4^+^ T cells exhibited significantly higher expression levels of the tested transcription factors, except for Foxp3, compared to their Foxp3^+^ counterparts (Figure 2E). The Foxp3^-^CD4^+^ T cell population can be further classified based on cytokine production patterns into subpopulations, such as IFN-γ^+^ (IFN-γ single positive; SP), IL-10^+^ (IL-10 SP), and IFN-γ^+^IL-10^+^ (double positive; DP). Among these, the IL-10 SP and DP subpopulations are classified as Tr1 cells [17, 20]. HK-C60 administration significantly increased the percentages of all three subpopulations within the Foxp3^-^CD4^+^ T cells. Notably, the most pronounced changes were observed in the DP subpopulation, followed by the IL-10 SP subpopulation (Figures 2F, G). Thus, HK-C60 administration enhances the Foxp3^-^CD4^+^ T cell population, particularly Tr1 cells, in the intestinal environment.

### C60 enhances IL-10 production and antigen-presenting ability in intestinal DCs

Our previous report demonstrated that C60 modulates intestinal DC function, characterized by increased cytokine production and antigen-dependent T cell activation [10, 11]. To explore how C60 influences intestinal DC function to promote IL-10^+^CD4^+^ T cell differentiation, we conducted RNA microarray analysis of intestinal DCs. Mice were administered saline or HK-C60 daily for 14 days, after which small intestinal PPs were excised, and cells were isolated. CD11c^+^ DCs were sorted from PP-derived cells, and total RNA was subjected to microarray analysis (Figure 3A). A volcano plot revealed that 450 genes were significantly upregulated, while 265 were downregulated in PP DCs from HK-C60-treated mice compared to controls (Figure 3B). To understand the functional impact of these changes, pathway analysis was performed. This analysis identified 543 upregulated pathways in C60-exposed intestinal DCs, with only 17 pathways being downregulated (Figure 3C). Notably, eight upregulated pathways were associated with IL-10 signaling, phagocytosis and micropinocytosis, and antigen presentation (Figure 3D). Heatmaps of extracted cytokine and antigen-presenting/co-stimulatory molecule genes showed significant differences in expression between control and C60-exposed PP DCs. Interestingly, HK-C60 administration markedly increased gene expression of both pro-inflammatory cytokines and the anti-inflammatory cytokine IL-10 in PP DCs compared to controls (Figure 3E). Additionally, genes encoding CD80, CD86, and MHC class II were upregulated in response to C60 (Figure 3F). These gene expression changes were validated at the protein level by flow cytometry. IL-10 production was significantly increased in DCs from the PP, sLP, and cLP of HK-C60-administered mice compared to controls (Figure 3G, H). Similarly, CD80, CD86, and MHC class II protein expression levels were all elevated in PP DCs following HK-C60 administration (Figure 3I, J).

**Figure 3.**
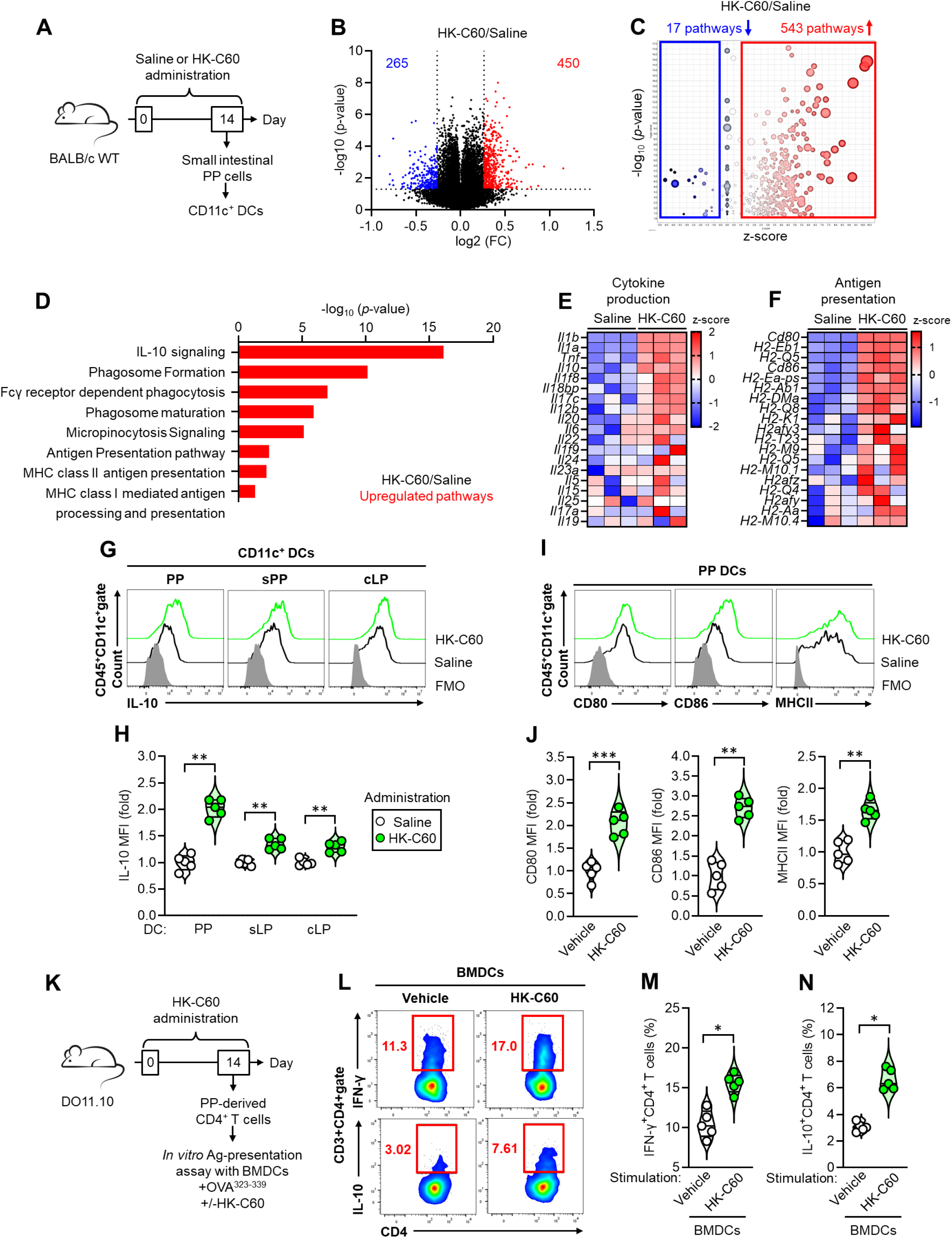
C60 enhances DC function accompanied by increased IL-10 production and antigen presenting ability. A) Experimental design of functional characterization of intestinal DCs. The mice received i.g. administration of saline or HK-C60 for 14 days. DCs were isolated from small intestinal PP-derived cells and subjected to RNA microarray or flow cytometry analysis. Total RNA was isolated from DCs and subjected to micro array analysis (n=3 in each group). B) Volcano plot of gene expressions. The red area indicated upregulated genes (FC ≥ 1.2) and blue area indicated downregulated gens (FC ≤ 0.75) with significant change (*p* < 0.05) in HK-C60-administered group compared to control group. C) Comprehensive pathway analysis following gene expression profile. D) Featured upregulated pathway in HK-C60-administered group. E-F) Heatmaps of cytokine production (E) and antigen presentation (F) associated gene expressions, respectively. G-H) Representative histogram (G) and cumulative MFI values (fold change) (H) of IL-10 production in intestinal DCs. I-J) Representative histogram (I) and cumulative MFI values (fold change) (J) of CD80, CD86 and HMC class II expressions in PP DCs. K) Experimental design of *in vitro* antigen presentation assay. Splenic CD4^+^ T cells were isolated from PP-derived cells of HK-C60-administered DO11.10 mice (14 days). The CD4^+^ T cells were co-cultured with BMDCs prepared from BALB/c WT mice in the presence of OVA_323-339_ peptide. The cultures were further treated with vehicle or HK-C60. After incubation at 37°C for 72 h, the cytokine production in CD4^+^ T cells was analyzed by flow cytometry. L) Representative plots of IFN-γ^+^ or IL-10^+^ CD4^+^ T cells. M-N) Cumulative percentages of IFN-γ^+^CD4^+^T cells (M) and IL-10^+^ CD4^+^ T cells (N) were shown, respectively. The cumulative data are shown as mean ± SEM values of five samples from two independent experiments. All MFI values are represented as fold changes (the average value of control group was used for base=1). Student t-test was used to analyze data for significance. \**p* < 0.01 and \*\**p* < 0.001, ns is not significant.

To assess how these functional modifications in DCs influence adaptive immunity, we performed an *in vitro* antigen presentation assay. DO11.10 mice, which harbor CD4^+^ T cells expressing a T cell receptor (TCR) specific for the ovalubmin (OVA) peptide antigen, were administered with HK-C60 for 14 days. CD4^+^ T cells were isolated from small intestinal PPs and co-cultured with bone marrow-derived dendritic cells (BMDCs) from wild-type (WT) BALB/c mice in the presence of OVA_323-339_ peptide. Cultures were treated with either saline or HK-C60. Flow cytometry analysis revealed that HK-C60 stimulation of BMDCs increased the frequency of IFN-γ^+^CD4^+^ T (Th1) cells, consistent with our previous findings [10, 11]. Notably, HK-C60 stimulation also significantly increased IL-10^+^CD4^+^ T cell frequencies compared to control treatment (Figure 3K-N). Importantly, HK-C60 alone did not promote effector CD4^+^ T cell differentiation upon TCR stimulation in the absence of DCs [11]. These results indicate that the functional modification of DCs by C60 is essential for the observed increases in effector CD4^+^ T cells. Specifically, enhanced IL-10 production in C60-stimulated DCs likely plays a key role in promoting IL-10^+^CD4^+^ T cell differentiation.

### C60 stimulated DCs induces IL-10^+^Tr1 cell differentiation from pro-inflammatory Th1cell

Previous studies have documented that IL-10^+^ DCs, referred to as DC10, play a crucial role in Tr1 generation by combining enhanced IL-10 production with increased antigen-presenting capacity [21, 22]. This raises the question of whether functionally modified intestinal DCs, stimulated by C60, promote IL-10^+^ Tr1 cell differentiation. Specifically, we investigated the mechanism through which IFN-γ^+^IL-10^+^ Tr1 cells differentiate from pro-inflammatory Th1 cells [23]. To address this, we developed a co-culture system to assess the contribution of C60-stimulated DCs in IL-10^+^ Tr1 cell generation. Splenic CD4^+^ T cells from DO11.10 mice were first cultured with IFN-γ, IL-12, and IL-2 in the presence of TCR stimulation to induce Th1 cell differentiation. The differentiated Th1 cells were then cultured with or without BMDCs in the presence of the OVA_323-339_ peptide. These cultures were treated with either vehicle or HK-C60, and Tr1 cells, identified as Foxp3^-^CD4^+^ populations, were further classified into subpopulations based on their cytokine production profiles (Figure 4A). In the absence of DCs, no IL-10-producing populations were observed within the Foxp3^-^CD4^+^ T cells, and equivalent Th1 cell populations were maintained in cultures treated with either vehicle or HK-C60. However, the presence of DCs, particularly those stimulated with HK-C60, significantly increased the percentages of IL-10^+^ and IFN-γ^+^IL-10^+^ populations, which are classified as Tr1 cells. Notably, the effect of HK-C60-stimulated DCs was more pronounced in promoting IFN-γ^+^IL-10^+^ Tr1 cell differentiation compared to that of IL-10^+^ Tr1 cell. By contrast, DCs cultured without HK-C60 stimulation induced only minimal increases in IL-10^+^ and IFN-γ^+^IL-10^+^ Tr1 cell populations (Figure 4B, C). These findings suggest that the functional modifications of DCs induced by C60 stimulation facilitate the differentiation of IL-10^+^ Tr1 cells from Th1 cells. This mechanism likely contributes to the reduction of excessive pro-inflammatory Th1 cells and the establishment of an anti-inflammatory environment.

**Figure 4.**
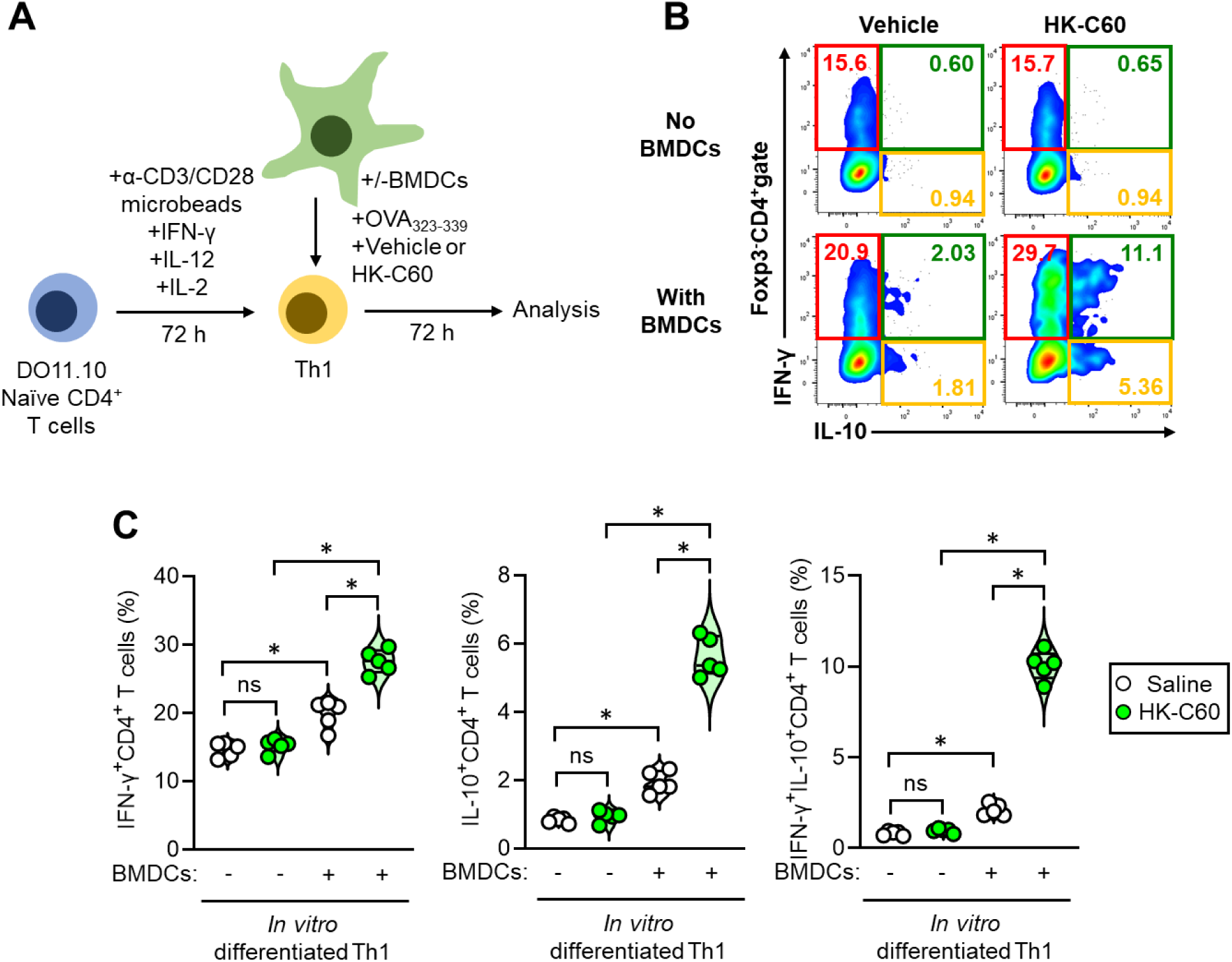
Functional modification of DCs by C60 promotes the IFN-γ^+^IL-10^+^Tr1 cell differentiation from pro-inflammatory IFN-γ^+^CD4^+^ T (Th1) cells. A) Experimental design of *in vitro* Tr1 cell differentiation assay. Splenic CD4^+^ T cells were isolated from DO11.10 mice and pre-cultured in the presence of anti-CD3/CD28 microbeads, IFN-γ, IL-12 and IL-2 at 37°C for 72 h to induce differentiation of IFN-γ^+^CD4^+^ T (Th1) cells. BMDCS were prepared from BALB/c WT mice. The precultured Th1 cells were further cultured with or without BMDCs in the presence of OVA_323-339_ peptide. Additionally, the cultures were treated with vehicle or HK-C60. After incubation at 37°C for 72 h, the cytokine production in Foxp3^-^ CD4^+^ T cells was analyzed by flow cytometry. B-C) Representative plots (B) and cumulative percentages (C) of IFN-γ^+^ (SP), IL-10^+^ (SP) and IFN-γ^+^IL-10^+^ (DP) Foxp3^-^CD4^+^ T (Tr1) cells. The cumulative data are shown as mean ± SEM values of five samples from two independent experiments. One-way ANOVA was used to analyze data for significance. \**p* < 0.001, ns is not significant.

### C60 administration protects against DSS-induced colitis accompanied by increased Tr1 cells

To evaluate the anti-inflammatory effects of C60 in a physiological setting, we employed the DSS-induced colitis model. Mice were administered saline or HK-C60 for 14 days. During the final 5 days, mice in the colitis group were given 5% DSS in their drinking water, while the control group received regular water. Body weight (BW) was monitored daily, and samples were collected at the end of the study for histological and immunological analyses (Figure 5A). In the saline-administered group, colitis induced by DSS caused significant, time-dependent BW loss. However, BW reduction was significantly mitigated in the HK-C60-administered group compared to the saline-administered group (Figure 5B). Similarly, DSS-induced colitis markedly shortened colon length in saline-administered mice, while this effect was milder in HK-C60-administered mice (Figure 5C, D). Histological analysis revealed that HK-C60 administration attenuated colon inflammation in DSS-induced colitis. The severity of inflammation was substantially reduced in the HK-C60-treated group compared to saline-treated mice (Figure 5E).

**Figure 5.**
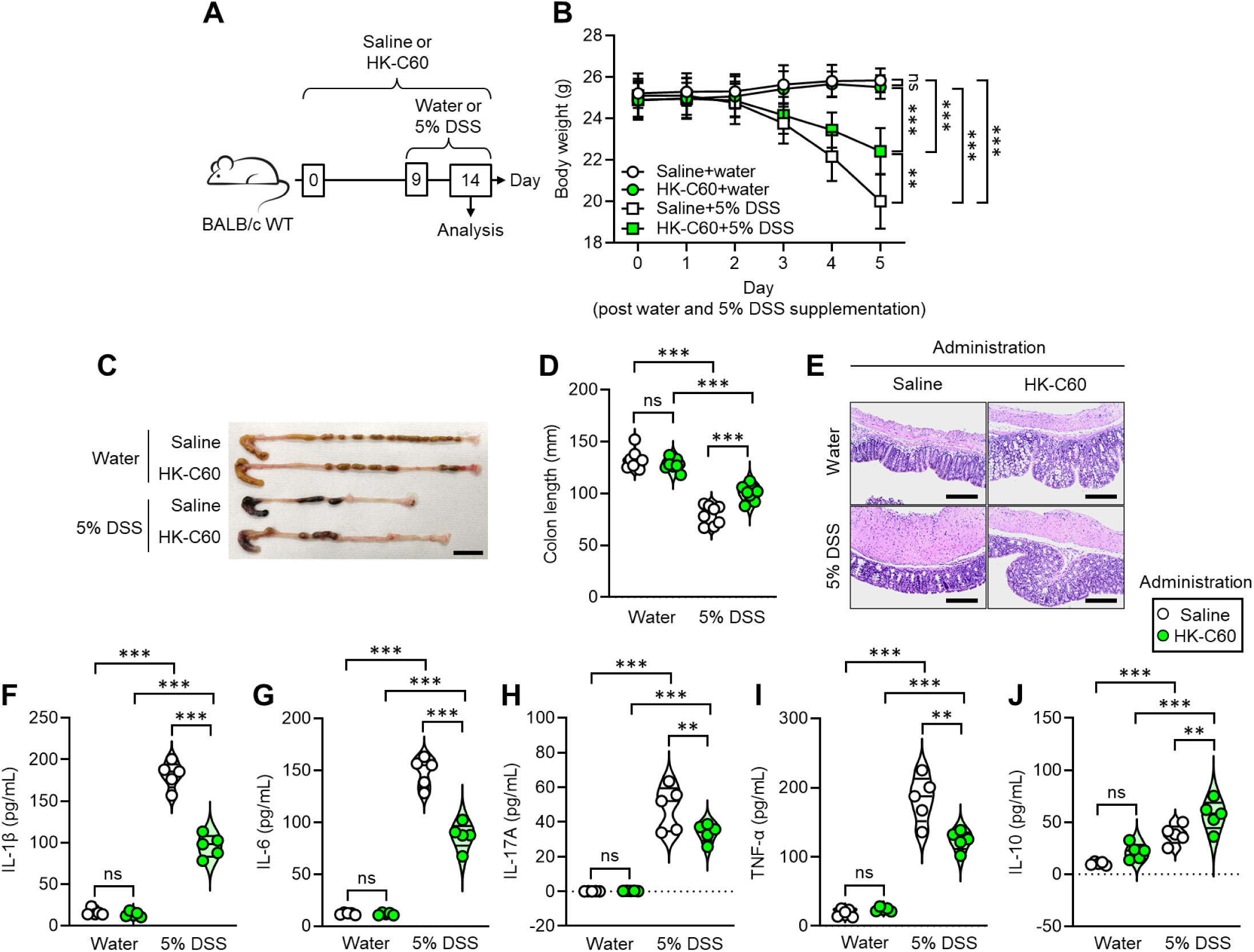
C60 administration attenuates inflammation in DSS-induced colitis. A) Experimental design of DSS-induced colitis. The WT mice received i.g. administration of saline or HK-C60 for 14 days. In the last 5 days, the mice were supplied with normal water or 5% DSS containing water. During this period, BW was measured in each mouse every day. In the last day of the experiment schedule, the mice were sacrificed and the intestinal tissues were subjected to analyses. B) The transition of BW. C) Representative pictures of appendix and colon. Bar=10 mm. D) Cumulative lengths of colon. E) Representative images of H-E staining of colon. Bar=100 µm. F-J) Measurement of colon cytokine productions. The colon section was homogenized, then supernatant was collected for cytokine ELISA. The concentrations of IL-1β (F), IL-6 (G), IL-17A (H), TNF-α (I) and IL-10 (J) were shown. The cumulative data are shown as mean ± SEM values of five samples from two independent experiments. One-way ANOVA was used to analyze data for significance. \**p* < 0.05, \*\**p* < 0.05 and \*\*\**p* < 0.001, ns is not significant.

To further assess inflammation, colon cytokine levels were analyzed. DSS-induced colitis significantly increased pro-inflammatory cytokines, including TNF, IL-1β, IL-6, and others, in the colon. Notably, HK-C60 administration significantly reduced the production of these cytokines compared to control treatment (Figure 5F-I). Intriguingly, IL-10 levels, known to counterbalance pro-inflammatory cytokines, were also markedly elevated in colitis tissues. This increase was more pronounced in HK-C60-treated mice compared to controls (Figure 5J).

Flow cytometric analysis showed that the percentage of IL-10^+^ Tr1 cells was already increased in the cLP under basal conditions (non-colitis) following HK-C60 administration, similar to the results observed in small intestinal PPs, as presented in Figure 2 (Figure 6A, top, and 6B). Under colitis conditions, the percentage of IL-10^+^ Tr1 cells increased significantly in both saline- and HK-C60-treated groups compared to the basal condition. This change was particularly pronounced in the IFN-γ^+^IL-10^+^ Tr1 cell subpopulation in HK-C60-treated mice, while the overall frequency of IL-10^+^ Tr1 cells was comparable between the two treatment groups (Figure 6A, bottom, and 6B).

**Figure 6.**
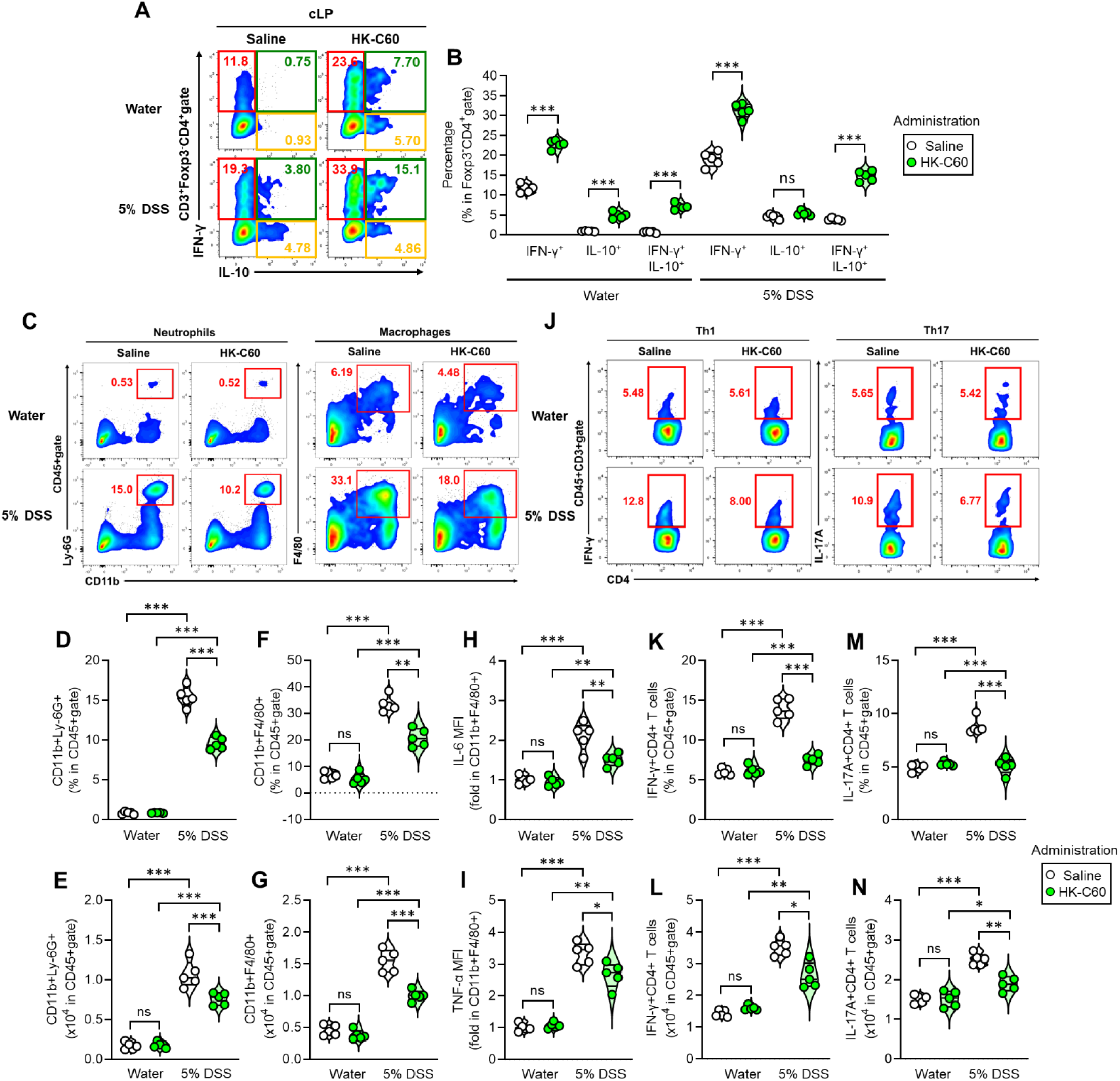
Increased Tr1 cell differentiation reduces inflammatory leukocytes in the intestine of C60-administered mice in DSS-induced colitis. The mice received i.g. administration of saline or HK-C60 for 14 days. In the last 5 days, the mice were treated with water or 5% DSS. The mice were sacrificed in the end of experiment, and intestinal immune profiling was performed by flow cytometry analysis. A-B) Cytokine production of Tr1 cells in cLP. Representative plots (A) and cumulative percentages (B) of IFN-γ^+^ (SP), IL-10^+^ (SP) and IFN-γ^+^IL-10^+^ (DP) populations of Foxp3^-^Tr1 cells are shown. C-G) Infiltrated myeloid cells in cLP. Representative plots (C), percentages (D, F) and cell numbers (E, G) of neutrophils (CD11b^+^Ly-6G^+^) and macrophages (CD11b^+^F4/80^+^) are shown. H-I) IL-6 production (H) and TNF-α production (I) in cLP macrophages. J-N) Infiltrated inflammatory T cells in cLP. Representative plots (J), percentages (K, M) and cell numbers (L, N) of IFN-γ^+^CD4^+^ T (Th1) cells and IL-17A^+^CD4^+^ T (Th17) cells are shown. The cumulative data are shown as mean ± SEM values of five samples from two independent experiments. All MFI values are represented as fold changes (the average value of control was used for base=1). One-way ANOVA was used to analyze data for significance. \**p* < 0.05, \*\**p* < 0.05 and \*\*\**p* < 0.001, ns is not significant.

Colon inflammation was further assessed by examining inflammatory immune cell infiltration into the cLP. Both neutrophils and macrophages, typically scarce in non-inflammatory conditions, were significantly increased in colitis. However, their infiltration was markedly reduced by HK-C60 administration (Figure 6C-G). Moreover, pro-inflammatory cytokine production from these myeloid cells was significantly decreased in the HK-C60-administered group compared to saline-administered colitis mice (Figure 6H, I). Inflammatory CD4^+^ T cells, including IFN-γ^+^CD4^+^ T (Th1) cells and IL-17A^+^CD4^+^ T (Th17) cells, also showed significant accumulation in colitis-affected cLP. These populations were significantly reduced in HK-C60-treated colitis mice compared to saline-treated controls (Figure 6J-N). Thus, HK-C60 administration confers protection against DSS-induced colitis by mitigating colon inflammation. This protective effect is associated with a pronounced increase in Tr1 cells and a reduction in pro-inflammatory immune cell infiltration and activity.

### Intestinal CD4^+^ T cells of C60-administered mice suppress inflammatory response of immune cells

We tested whether CD4^+^ T cells originating from C60-administered mice could suppress inflammatory responses of other T cells and macrophages using an *in vitro* co-culture system. Three different resources were prepared from distinct settings. As anti-inflammatory effector cells, CD4^+^ T cells were isolated from the small intestinal PPs of saline- or HK-C60-administered mice. The target CD4^+^ T cells with pro-inflammatory cytokine production were isolated from the small intestinal PPs of DSS-induced colitis mice. For inflammatory macrophages as targets, TPMs were prepared from naïve mice and then primed with LPS. The target inflammatory CD4^+^ T cells or macrophages were co-cultured with CD4^+^ T cells obtained from either saline- or HK-C60-administered mice, and the inflammatory responses of the target cells were analyzed by flow cytometry (Figure 7A). The PP-derived CD4^+^ T cells from HK-C60-administered mice significantly suppressed IFN-γ and IL-17A production in target intestinal CD4^+^ T cells compared to effector cells derived from saline-administered mice, even under basal conditions (derived from water-treated mice). This suppressive effect was more pronounced against colitis mouse-derived CD4^+^ T cells, which exhibited substantially increased IFN-γ and IL-17A production (Figure 7B, C). The TNF-α production in LPS-primed macrophages was also significantly suppressed by PP-derived CD4^+^ T cells from HK-C60-administered mice compared to control cells (Figure 7D). *In vitro* IL-10 neutralization revealed that this suppressive effect predominantly depended on the increased IL-10 secretion from PP-derived CD4^+^ T cells of HK-C60-administered mice. Cultures treated with an anti-IL-10 mAb showed abolished anti-inflammatory effects of the intestinal IL-10-producing CD4^+^ T cells originating from HK-C60-administered mice (Figure 7E-G). Thus, the expanded intestinal IL-10^+^ CD4+ T cells induced by C60 administration have the capability to suppress the inflammatory responses of other immune cells in an IL-10-dependent manner.

**Figure 7.**
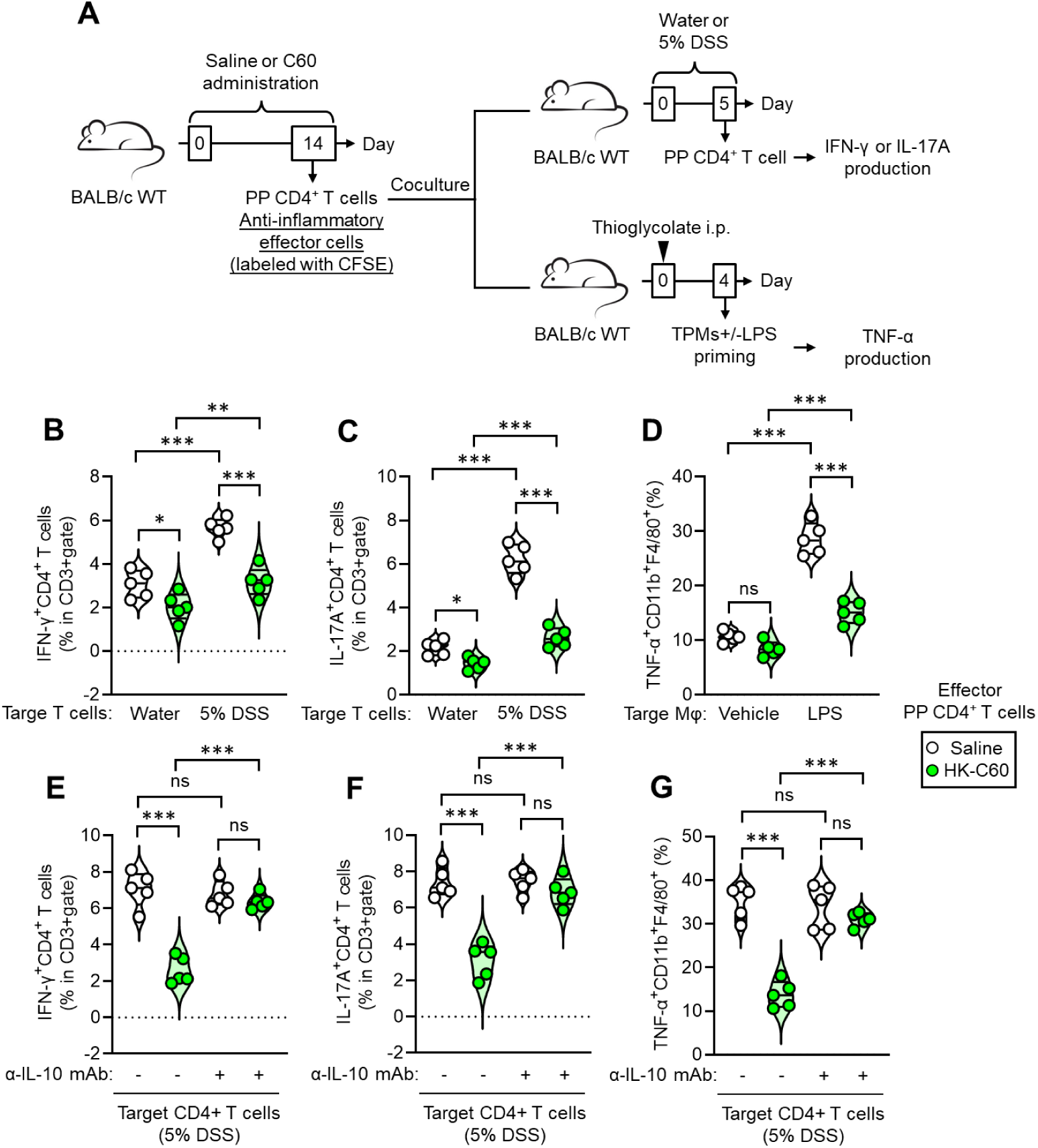
Intestinal CD4^+^ T cells in C60-administered mice possess IL-10-dependent suppressive effect against inflammatory immune cells. A) Experimental design of *in vitro* co-culture for suppression assay. The effector CD4^+^ T cells were isolated from small intestinal PPs of WT mice received saline or HK-C60 administration for 14 days. The administration was performed by following the method described in materials and methods. The effector CD4^+^ T cells were labeled with CFSE before use to distinguish them from target cells. The target inflammatory CD4^+^ T cells were isolated from the small intestinal PPs of WT mice applied water or 5% DSS containing water for 5 days. The TPMs were prepared from naïve WT mice following the protocol described in materials and methods. The effector and target CD4^+^ T cells were mixed and co-cultured at 37°C for 24 h, then the frequency of IFN-γ or IL-17A producing cells in target CD4^+^ T cell population was analyzed by flow cytometry. The TPMs were first primed with LPS at 37°C for 6 h followed by co-cultured with effector CD4^+^ T cells. The frequency of TNF-α producing cells were analyzed by flow cytometry. Alternatively, some cultures co-cultures were treated with isotype (-) or anti-IL-10 mAb (+) for neutralization. B-D) Cumulative percentages of IFN-γ^+^CD4^+^ T (Th1) cells (B), IL-17A^+^CD4^+^ T (Th17) cells (C) and TNF-α^+^ macrophages (CD11b^+^F4/80^+^) (D) in suppression assays were shown, respectively. E-G) Cumulative percentages of IFN-γ^+^CD4^+^ T (Th1) cells (E), IL-17A^+^CD4^+^ T (Th17) cells (F) and TNF-α^+^ macrophages (CD11b^+^F4/80^+^) (G) in suppression assays with or without IL-10 neutralization were shown, respectively. All CD4^+^T cells were stimulated with anti-CD3/CD28 microbeads (at the ratio=1:1). The cumulative data were shown as mean +/− SEM of five samples in two independent experiments. One-way ANOVA was used to analyze data for significant differences. Values of \**p* < 0.05, \*\**p* < 0.01 and \*\*\**p* < 0.001 were regarded as significant. ns; not significant.

## Discussion

The immunomodulatory effects of probiotic lactic acid bacteria (LAB) are well-recognized, with anti-inflammatory properties being a significant area of interest. As described in this report, increased IL-10-producing CD4^+^ T cells play a predominant role in probiotics-mediated anti-inflammatory responses. Regarding anti-inflammatory T cell behavior, most studies primarily focus on Treg as the subset exhibiting this characteristic [16]. However, Tr1 cells are also capable of producing IL-10 [17–23], and our result revealed that the IL-10-producing capacity is higher than that of Treg. This evidence supports the promising potential of strategies aimed at increasing Tr1 cell generation to suppress intestinal inflammatory diseases and immunological abnormalities.

However, the mechanisms to reliably achieve this remain largely unexplored. LAB-based probiotics represent one scientifically credible approach for this purpose [20]. Our study highlights that the probiotic strain C60 preferentially promotes Tr1 cell generation rather than Treg differentiation. This finding underscores the potential of C60 as a probiotic strain that specifically enhances Tr1 cell-mediated immune modulation. While our results do not discount C60’s role in promoting Treg differentiation, the predominant effect is observed in Tr1 cell population changes. Additionally, our data revealed that functional modifications in DCs are the primary mechanism underlying the increase in IL-10^+^CD4^+^ T cells in the intestine. DCs exposed to C60 demonstrated significantly increased IL-10 production alongside upregulated pro-inflammatory cytokine productions and enhanced antigen-presenting capabilities [10, 11]. These functional modifications are unique to C60, as shown by comparisons with other LAB strains [11].

In the context of antigen-dependent anti-inflammatory T cell differentiation, our findings show that C60-stimulated DCs are capable of inducing Tr1 cell differentiation, specifically in IFN-γ^+^IL-10^+^ populations, which may originate from pro-inflammatory IFN-γ^+^CD4^+^ T (Th1) cells. This novel finding suggests that DCs functionally modified by C60 facilitate the differentiation of unique anti-inflammatory T cells from Th1 cells. This mechanism likely contributes to creating an anti-inflammatory immune environment by reducing excessive Th1 cells [24], known inflammatory inducers, thereby mitigating intestinal inflammation risks. Nonetheless, careful consideration must be given to maintaining the balance between pro-inflammatory and anti-inflammatory T cells in the intestinal environment. Sustaining sufficient Th1 cell populations is crucial for a well-conditioned intestinal immune environment. Notably, C60 probiotics are expected to support this balance, as documented in our previous study [10]. Thus, we emphasize that C60 offers comprehensive immunomodulatory functions, enhancing both Th1 cell and Tr1 cell activities to maintain intestinal health.

The mechanisms by which C60-stimulated DCs influence the differentiation of Tr1 cell from Th1 cell remain to be fully elucidated. In the context of antigen-dependent Tr1 cell generation, previous studies have reported that IL-10^+^DCs (DC10) play a crucial role [21, 22]. Both TCR-dependent signaling and proximally enriched IL-10 appear to be indispensable for Tr1 cell differentiation. The DC phenotype induced by C60 closely resembles that of DC10, characterized by enhanced antigen presentation through upregulated expression of associated molecules and increased IL-10 production. Neutralizing IL-10 significantly reduced IL-10^+^CD4^+^ T cell differentiation in our in vitro co-culture system, highlighting the importance of proximal IL-10 in promoting these anti-inflammatory lymphocytes differentiation upon antigen-dependent stimulation. However, the precise mechanisms through which the IL-10 network influences CD4^+^ T cells during IL-10^+^ differentiation remain unclear. Hypothetical pathways suggest that DC-derived IL-10, acting through the IL-10 receptor (IL-10R) on CD4^+^ T cells, may play a crucial role in upregulating gene expression associated with IL-10^+^CD4^+^ T cell differentiation via signal transducer and activator of transcription 3 (STAT3) activation [25, 26]. Addressing these pathways will be a focus of future research.

In a DSS-induced colitis model, C60 administration significantly suppressed intestinal inflammation by increasing IL-10^+^CD4^+^ T cells, predominantly composed of Tr1 cells rather than Tregs. Remarkably, C60 reduced the infiltration of inflammatory immune cells into the cLP and suppressed pro-inflammatory cytokine production in colon tissues, while increasing IL-10 production. This finding suggests that C60 has potential applications beyond intestinal inflammation, warranting further exploration of its effects in other inflammatory diseases.

In conclusion, our results provide strong evidence for the immunobiological contributions of C60 in probiotic-based approaches. As a probiotic strain, C60 holds promise for comprehensive immunomodulatory effects that support intestinal health. While further studies are needed to clarify underlying mechanisms, C60 demonstrates strong potential as a therapeutic tool for maintaining intestinal immune homeostasis.

## Acknowledgment

We thank the National Agriculture and Food Research Organization (NARO) for providing *Lactococcus lactis* subsp. *cremoris* C60.

## Author Contributions

SS, NK, AO, and TM performed the experiments. SS conducted data analysis and finalized the figures. SS and NMT designed the study and established the methodologies. The study was conceptualized by SS and NMT. NMT provided resources, supervised the study, and was responsible for project administration. SS drafted the original manuscript, and SS and NMT finalized it. All authors participated in discussions.

