## Supplemental Materials for "*Lactococcus lactis* subsp. *cremoris* C60 creates a type I regulatory T cell-dominant anti-inflammatory intestinal environment through functional modification of dendritic cells"

Address correspondence to: Noriko M. Tsuji, Ph.D., Division of Immune Homeostasis, Department of Pathology and Microbiology, Nihon University School of Medicine, Itabashi, Tokyo 1738610, Japan

---

**Supplemental Table 1. Primer sequences for real-time qPCR**

| Target gene | Forward (5'-3') | Reverse (5'-3') |
|---|---|---|
| *Foxp3* | CCCATCCCCAGGAGTCTTG | ACCATGACTAGGGGCACTGTA |
| *Eomes* | GCGCATGTTTCCTTTCTTGAG | GGTCGGCCAGAACCACTTC |
| *c-Maf* | GGAGACCGACCGCATCATC | TCATCCAGTAGTAGTCTTCCAGG |
| *Batf* | CTGGCAAACAGGACTCATCTG | GGGTGTCGGCTTTCTGTGTC |
| *Blimp1* | TTCTCTTGGAAAAACGTGTGGG | GGAGCCGGAGCTAGACTTG |
| *Irf1* | ATGCCAATCACTCGAATGCG | TTGTATCGGCCTGTGTGAATG |
| *Irf4* | TCCGACAGTGGTTGATCGAC | CCTCACGATTGTAGTCCTGCTT |
| *Gapdh* | AGGTCGGTGTGAACGGATTTG | TGTAGACCATGTAGTTGAGGTCA |

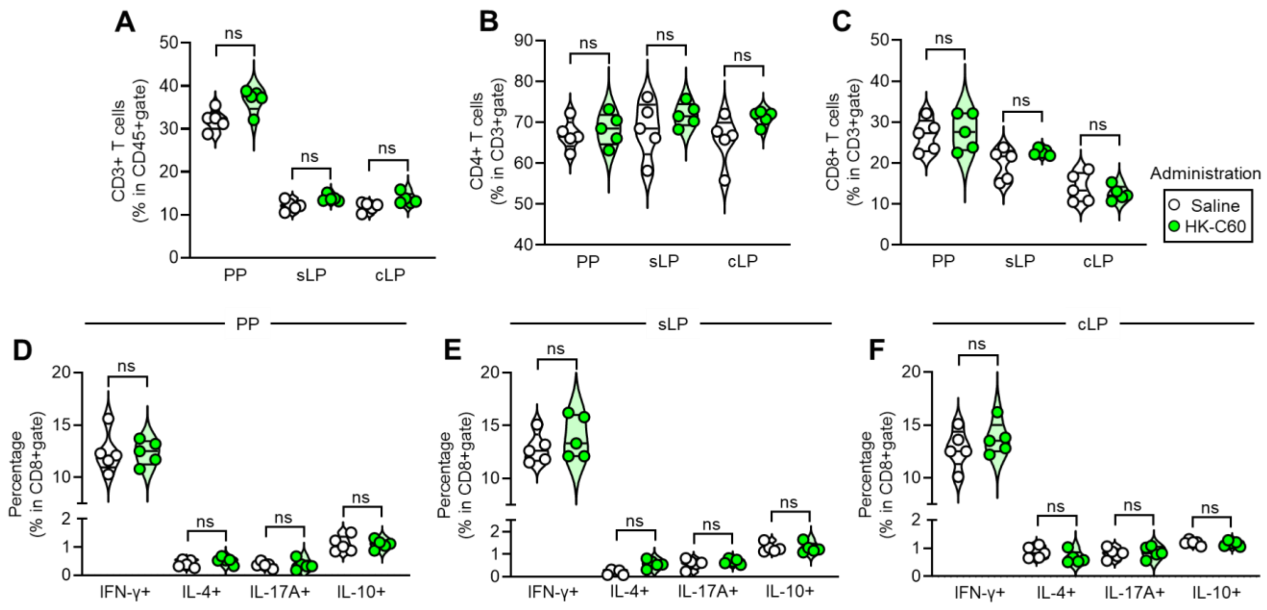


**Supplemental Figure 1. T cell immunophenotyping of intestinal environment.** The mice received saline or HK-C60 administration for 14 days, then the cells were isolated from each intestinal tissue. The T cell phenotypes were classified by following cytokine producing patters in flow cytometry analysis. The graphs show the percentages following the presented categories: A) Total $CD3^+$ T cells, B) Total $CD4^+$ T cells, C) Total $CD8^+$ T cells in PP, sLP and cLP. Effector $CD8^+$ T cells in PP (D), sLP (E) and cLP (F), respectively. The cumulative data are shown as mean ± SEM values of five samples from two independent experiments. Student t-test was used to analyze data for significance. ns is not significant.

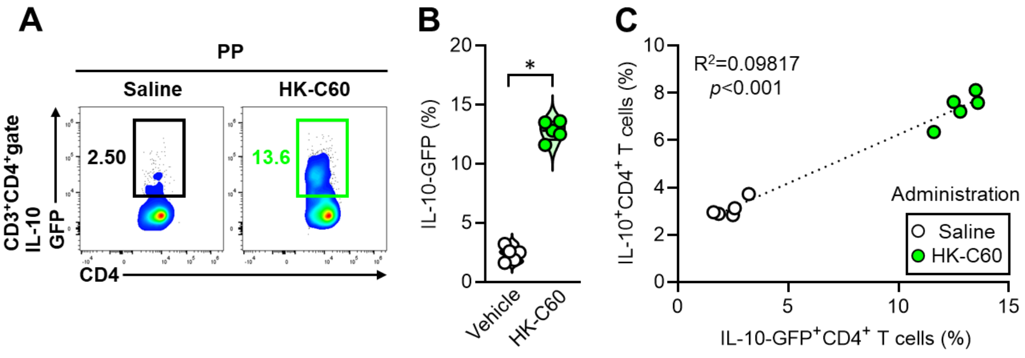


**Supplemental Figure 2. GFP expression in intestinal $CD4^+$ T cells of IL-10-GFP reporter mice**

IL-10-GFP mice received saline or HK-C60 administration for 14 days, then the small intestinal PP-derived cells were isolated to analyze GFP signals theoretically corresponding to the expression level of IL-10. At the meantime, IL-10 was directly stained in the cells after stimulation following protocol described in materials and method section, then analyzed by flow cytometry. A) Representative plots of IL-10-$GFP^+CD4^+$ T cells. B) Cumulative percentage values of IL-10-$GFP^+CD4^+$ T cells. C) Correlation of percentages between IL-10-$GFP^+$CD4+ T cells and IL-$10^+CD4^+$ T cells in the identical samples. The cumulative data are shown as mean ± SEM values of five samples from two independent experiments. Student t-test was used to analyze data for significance. Value of *$p<0.001$ was regarded as significant. Linear regression analysis was performed by using GraphPad Prism 10.0.
